# Patients with different cancer types are stratified by CBC data

**DOI:** 10.1101/796078

**Authors:** Michael G. Sadovsky, Alena A. Feller, Elena A. Martynova, Denis V. Chernyaev, Edward V. Semenov, Eugene V. Slepov, Ruslan A. Zukov

**Affiliations:** Siberian federal university, Center for genomic research, Krasnoyarsk, Russia; Institute of computational modelling of SB RAS, Krasnoyarsk, Russia; V. F. Voino-Yasenetsky Krasnoyarsk state medical university, Krasnoyarsk, Russia; A. I. Křyzhanovski Krasnoyarsk Regional Clinical Oncology Hospital, Krasnoyarsk, Russia

## Abstract

Searching for informative indices indicating cancer type and cause of the disease is of great importance. Here we tried to identify those indices from the data of complete blood analysis, only. We studied the inhomogeneity in the mutual distribution of a number of oncology patients with various types of tumors in the space of qualitative data provided by the complete blood count. The patients with oncology in hematology system were excluded. Ultimate goal is to reveal the relation between such inhomogeneity issues and the cause of a disease. We used the database on complete blood count comprising oncology patients with various causes of tumor development. The analysis has been carried out both by linear (*K*-means) and non-linear (elastic map technique) methods. No linear clustering has been found. On the contrary, elastic map technique yields stable clustering identifying not less than three clusters, in the set of patients. No relation of those clusters to sex or age of patients has been found. Four indices (namely, BAS, EOS, WBC and IG) exhibit no relation to the cluster structure, while all others do it. Thus, the patients are stratified according to their respond on the stress caused by cancer tumor. The data on complete blood count may be used for preliminary diagnostics of a tumor and its cause, for oncology patients. This type of analysis is cheap, standard and available at any medical organization.

## Introduction

Malignant neoplasm is among the leading factors of death rate, worldwide. Currently, a number of various techniques and methods are used to detect, study, trace and treat cancer tumors [1–3]; those methods range from highly advanced based on molecular genetics to classic data records. It should be also stressed that many up-to-date methods and techniques of diagnostics and treatment of cancer diseases are very expensive and complicated in practice [4]. Yet, the problem is far from a final resolution: there are quite a number of cancer types escaping from any up-to-date early diagnostics methods. *brca*1 and *tnbc* types of that latter are the typical examples. In particular, *brca*1 cancer type could be detected through molecular genetics tests, mainly, while *tnbc* is the breast cancer undetectable by standard techniques [5–7].

Hence, a development and implementation of various techniques and approaches to detect a malignant neoplasm, as well as to treat it effectively makes the core issue of an up-to-date medicine and medical biology sciences. The most advantageous methods contribute a lot both scientific oncology and cancer treatment; simultaneously, various data banks on cancer and related problems are currently available to researchers, and big data approach may bring a lot here [8]. Such data banks are full of hidden or conspired relations between the numerous records, figures and characters; a researcher may capitalize a lot from careful knowledge extraction and examination of such data banks.

Key idea here is a search for some inhomogeneities in the points representing the data, in multidimensional space, as well as newly identified interdependencies between the variables or their combinations, rather than a verification of *á priori* formulated hypotheses. Of course, one should seek the most informative or significant variables or their combinations yielding the inhomogeneities mentioned above; these former may be of high informative value both for a researcher, and a doctors. The idea is a following: firstly, a researcher identifies inhomogeneities in the data points distribution, and checks whether these clusters are distinguishable. At the second step an interplay between the composition of those clusters and the characteristics of a disease (or a patient) is investigated. This approach is very sounding for an investigation of data banks containing abundant clinical records collected in hospitals or other medical services.

The approach mentioned above might be applied to analysis of the data records representing some routine, widely spread, customized and highly widespread medical analysis data, thus bringing some new understanding on the problem. Here we analyzed the data bank on complete blood count supported by A. Křižanovski Krasnoyarsk Oncology Hospital, Krasnoyarsk, Russia. The dataset contains the records of the complete blood count of the patients suffering from various oncological diseases, excepting the blood system cancer, itself. Again, this analysis is mandatory in any medical practice, and its cost an availability alongside the long story of its implementation make the data absolutely reliable and consistent, nationwide.

The idea to collate the data records on complete blood count and oncological pathology is not absolutely new. Paper [9] shows the efficiency of complete blood count (CBC) to foresee and correct the treatment course of patients with oncology of two types: mammal cancer, and colon cancer. CBC data were used to trace quite specific treatment procedure of those pathologies that is intraoperative chemotherapy provided be automedia. A relation between CBC records and ovarian cancer is discussed in [10]. The paper shows the possibility to improve an early diagnosis of the disease through the complex analysis of CBC records. The relations between CBC, biochemical and immunological data obtained from the blood of patients with mammal cancer are presented in [11]. The authors believe that the significant intragroup scattering absence may reflect the stability of the metabolism processes and general sustainability of a patient. Differential diagnostics of carcinoid tumors of gastrointestinal tract including CBC data is discussed in [12] and CBC is shown to be efficient tool to differential diagnostics of some types of cancer of gastrointestinal tract.

CBC could be used as a prognostic marker of the metastasis probability in lymph nodes caused by endometrial cancer [13]. The patients came through complete cure course against the disease mentioned above could be reliable classified into the risk group vs. the group with minimal probability of metastasis. The classification has been carried out due to linear logistic regression; that latter revealed the hidden relation between the growth of neutrophil cells and lymph nodes damage. Similarly, paper [14] provides an estimation of the endometrial cancer probability based on CBC records. They used the characters of erythrocyte distribution width over the volume of these latter as the biomarkers related to inflammatory process. Also, these distribution parameters are prognostic very informative for patients with endometrial cancer. Again, the authors stress the easy-to-do and very low cost of CBC recording.

Paper [15] reports on the feasibility of CBC in colon cancer examination, for the purposes of early diagnostics. Some CBC figures indicate unambiguously the increased risk of coloteral cancer, that in turn results in the assignment of extra special examination for a patient. The approach is based on machine learning methods [16]; in such capacity, this paper resembles, to some extent, our approach and results. The correlation analysis between pre-surgery and post-surgery thrombocyte content associated with some surgery factors for patients with cervical cancer is reported in [13]. This paper claims that some figures of CBC could be used as a supplementary tool for pre-surgery diagnostics in cervical cancer patients. Finally, the papers [17, 18] strongly correspond to the approach we pursue here: it presents the studies of simple screening techniques including CBC to get a diagnosis of myeloid leukemia. The method stands behind a simple and easy-to-do technique for screening of myeloid leukemia. It was found that CBC figures, namely, the absolute figures of basophiles content effectively detects chronic myeloid leukemia. Key questions of the paper are whether it is possible to predict the out-put of some very expansive and complicated examinations over CBC data, and if yes then what is the accuracy of this prediction. It was found that there are some relations between various CBC figures that for sure predict the disease development.

Thus, the aim of our research is to identify the inhomogeneities in the distribution of the patients suffering from oncological tumors of different localization, origin and type, and to reveal the relations between some features of the diseases and the composition of clusters identified in the multidimensional data space. Each patient record is the string of figures obtained from the complete blood count, thus comprising a data set in multidimensional metric space. An interplay between the structure revealed in the space, and the peculiarities of the disease characteristics is the ultimate goal of this work.

## Materials and methods

We studied the database on CBC records obtained from patients with different oncology (excluding various types of leucemia), sex and age of a patient. All data have been obtained due to standard techniques, besides each record contained the information on cancer type (encoded within *International Statistical Classification of Diseases and Related Health Problems* codes). The blood samples were obtained from the patients of Křižanovski Krasnoyarsk Oncology Hospital. The database comprises 867 entries; following CBC parameters have been included into the database: BAS (basophils absolute abundance in peripheral blood), EOS (eosinophils absolute abundance in peripheral blood), HCT (haematocrit content), HGB (haemoglobin concentration in whole blood), IG (absolute abundance of immature granulocytes in peripheral blood), LYM (lymphocytes absolute abundance in peripheral blood), MCH (average haemoglobin content per an erythrocyte), MCHC (saturation level of erythrocytes with haemoglobin), MCV (erythrocyte average volume), MON (monocytes absolute abundance in peripheral blood), MPV (average volume of a thrombocyte), NEUT (neutrophiles absolute abundance in peripheral blood), P-LCR (portion of large thrombocytes), PCT (thrombocrit content), PDW (relative width of thrombocyte distribution over the volume), PLT (thrombocytes absolute abundance in peripheral blood), RBC (erythrocytes absolute abundance in peripheral blood), RDW-CV (variation of the relative width of thrombocyte distribution over the volume), RDW-SD (standard deviation of the relative width of thrombocyte distribution over the volume), WBC (leucocytes absolute abundance in peripheral blood) and erythrocyte sedimentation rate (ESR).

Data pretreatment includes a routine statistical analysis, namely

- for each variable the average and standard deviation have been determined;
- each variable has been tested against the normality of the distribution;
- finally, histogram has been developed for each variable, to evaluate the distribution pattern.

Principle component analysis (PCA) [19] has been implemented at the next stage, as well as the correlation matrix calculation. PCA has been applied to determine the efficient (linear) dimension of the data, and correlation matrix has been calculated to determine the couples of the variables with high linear constraint. That latter has been used to select the variables from CBC records to be eliminated from further analysis; the point is that numerous linear constraints may cause some bias in inner structure revealing.

Following couples of variables HCT-HGB (0.96), MCH-MCV (0.91), LYM-NEYT (0.96), MPV-PLCR (0.95), MPV-PDW (0.93), PLCR-PDW (0.94), PCT-PLT (0.96), RDWCV-RDWSD (0.85) have been fond to exhibit high and very high correlation coefficients. We applied the threshold of the variable exclusion from the further analysis equal to ≥ 0.85; hence, the variables MCV, NEUT, MPV, PDW, PLCR, PLT, RD-WSD were excluded from further analysis.

Hence, each patient is represented by the point in 21-dimensional metric space. Everywhere below we used Euclidean metrics to measure the distance in the space. Table 1 shows the abundances of neoplasms of various type located in patients that are included into the database. So, we sought for the inhomogeneities in the distribution of the patients enrolled into the database, in the 21-dimensional Euclidean space of CBC figures, only. Next, if such clusters are found, then what is their composition in terms of pathology, patient features or disease characteristics. Reciprocally, the dual question was also examined. We studied the pattern and level of homogeneity in the patients distribution in different clusters. In other words, it was rather important to learn to what extend a cluster comprises the patients with different features.

**Table 1:**
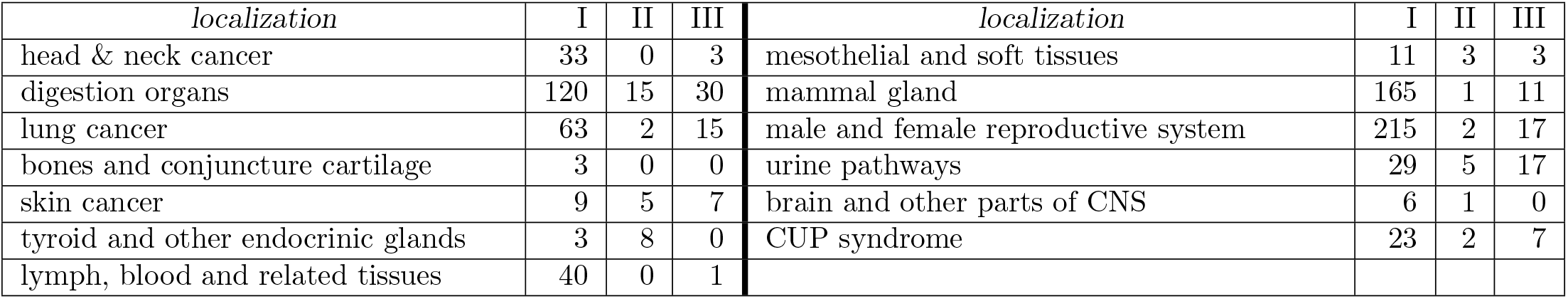
Localization and type of neoplasm. Here I stands for malignant neoplasm, II stands for benign tumor, and III stands for unknown type.

| *localization* | I | II | III | *localization* | I | II | III |
| --- | --- | --- | --- | --- | --- | --- | --- |
| head & neck cancer | 33 | 0 | 3 | mesothelial and soft tissues | 11 | 3 | 3 |
| digestion organs | 120 | 15 | 30 | mammal gland | 165 | 1 | 11 |
| lung cancer | 63 | 2 | 15 | male and female reproductive system | 215 | 2 | 17 |
| bones and conjuncture cartilage | 3 | 0 | 0 | urine pathways | 29 | 5 | 17 |
| skin cancer | 9 | 5 | 7 | brain and other parts of CNS | 6 | 1 | 0 |
| thyroid and other endocrinic glands | 3 | 8 | 0 | CUP syndrome | 23 | 2 | 7 |
| lymph, blood and related tissues | 40 | 0 | 1 |  |  |  |  |

### Clustering and visualization techniques

Let now briefly outline the clustering methods used in our study. We used two methods of clustering: linear and nonlinear ones. The former is *K*-means, and the latter is elastic map technique. *K*-means is well-described and comprehensively investigated classification method, see, e. g. [19] for details. Elastic maps technique is the powerful and advanced method to approximate the data points in multidimensional space by a manifold of low dimension; practically, they use two-dimensional manifolds. The trick is that one-dimensional manifold approximation is harder to do and more complicated both in implementation and interpretation. An idea standing behind the elastic map technique is to adjust the points in multidimensional space as precisely, as possible, while reaching the minimal (in some sense) transformations of the manifold. Elastic map technique provides the non-linear dimension reduction: indeed, it reduces the dimension of the data set to two.

Let now describe elastic map technique in more detail. The first step consists in well-known traditional approach that is PCA [19]. Geometrically, the first principal component is the direction at the original Euclidean space providing that the data are mostly scattered along it. So, one must determine the first and the second principal components and spread a plane on them as on axes. Next, each data point must be projected on the plane, and minimal square comprising all the projections must be determined. At the second step, each data point must be connected to its projection with (mathematical) spring; all the springs here are stipulated to be equal, in their elasticity. It means that all the springs have the same original length (equal to zero, to be exact), and the same elasticity. Moreover, the springs are supposed to be infinitely expandable and the linear elasticity is expected, for any extension length.

When all the points are connected to their projections with mathematical spring, one must change originally (infinitely) rigid plane (the square, to be exact) for elastic membrane. A membrane is supposed to be uniform and homogeneous: it has the same elasticity properties in any point, in in all directions. So, at the third step the system is released to reach the minimum of total deformation energy. That latter is a combination of elastic membrane deformation (expansion, torsion and bending), and the mathematical springs expansion energy.

As soon, as the equilibrium configuration is achieved, each data point must be redetermined over the jammed surface. Indeed, the orthogonal projection for each point must be found; geometrically, this new point image is located on the jammed surface at the closest distance from the original point. The elastic map is ready for use. To reveal cluster structure, one should transform back the jammed surface into a plane state. To do that, one must cut-off all the mathematical springs, so that the jammed surface becomes plane. This is the nonlinear transformation into the so called inner coordinates. Obviously, straightening the jammed surface results in a change of the location of the images of the data set points (these were orthogonal projections). The projected images located on a convex part of the map will get closer, as the map straightens; vice versa, the projected images located on a concave part of the map will get more distant. This is intuitively clear, since the first case means that the original points were “strong enough” to pull the elastic map to them; reciprocally, the second case means that the points were “weak enough” to keep the elastic map close to them.

Thus, the backward transformation straightening the map allows to visualize and cluster (in non-linear mode) the points located in multidimensional space. So, the final pattern depends on elasticity of the map and the strings. We used freely distributed software *VidaExpert* [20]. Practically, due to computational constraints, nobody implement elastic map in a software as a continuous object. On the contrary, elastic map is presented as polyhedron, and the points are connected to the nodes of that latter, not the exact projection image. There may be three types of elastic map, in dependence on the number of nodes in a square: rigid (usually 12 × 12), soft (16 × 16) and detailed (25 × 25); we used the soft configuration in our studies.

To visualize the newly obtained image points distribution over the map, one should introduce some local density clustering procedure. There are a number of this kind methods (see, e. g., [21] for some details), and we shall implement probably the simplest one based on local density function. To do that, one must supply each image point on the elastic map with Gaussian function

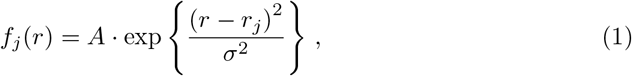

where *r_j_* is the coordinate vector (in inner coordinates) of *j*-th point. *r* is the radius originated at *r_j_, A* is the factor mainly equal to 1, and *σ* (looking similar to standard deviation in normal distribution) is the contrasting parameter: it determines the width of the “hat” covering a point. It should be stressed that function (1) is rotationally symmetric, for the sake of simplicity. Finally, the sum function

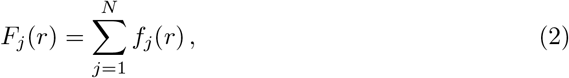

where *N* is the total number of points in the data set. Function (2) shows the local density function representing a cluster structure in the data set, if any.

## Results

To begin with, let’s consider PCA results and correlation analysis of the original data set. To determine the efficient dimension of the data set in linear approximation, we have carried out principal component analysis; in particular, we have started from eigenvalues calculation for the covariation matrix obtained from the data. Fig. 2, left, shows the variation of the eigenvalues, as the number of that latter grows up. The curve has a distinct break at the second eigenvalue; all other eigenvalues follow almost linear pattern of the decrease. Thus, the efficient space dimension in our data base is equal to one; nonetheless, one hardly could approximate the data with a one-dimensional linear manifold: other eigenvalues are not small enough to omit them. Practically, it means that the data are mainly dispersed along a line, but this distribution is rather wide and non-trivial.

Fig. 2, right, shows the distribution mentioned above. The figure shows the data distribution in special projection, where the first principal component is directed orthogonally to the figure plane. Thus, the axes on this figure represent the contribution of each variable into the distribution pattern.

We studied a classification of the patients in the database by *K*-means. No reasonable classification has been observed for 2 ≤ *K* ≤ 6; all the classifications were very unstable. The instability means that the greater part of the patients changed their class attribution, in a series of *K*-means runs. Hence, we have changed for elastic map technique. Fig. 1 show the pattern of clustering obtained by the soft (16 × 16) elastic map, with by default parameters of elasticity. Evidently, there are three or four clusters on the map, in dependence on the contrast parameter.

**Fig 1.**
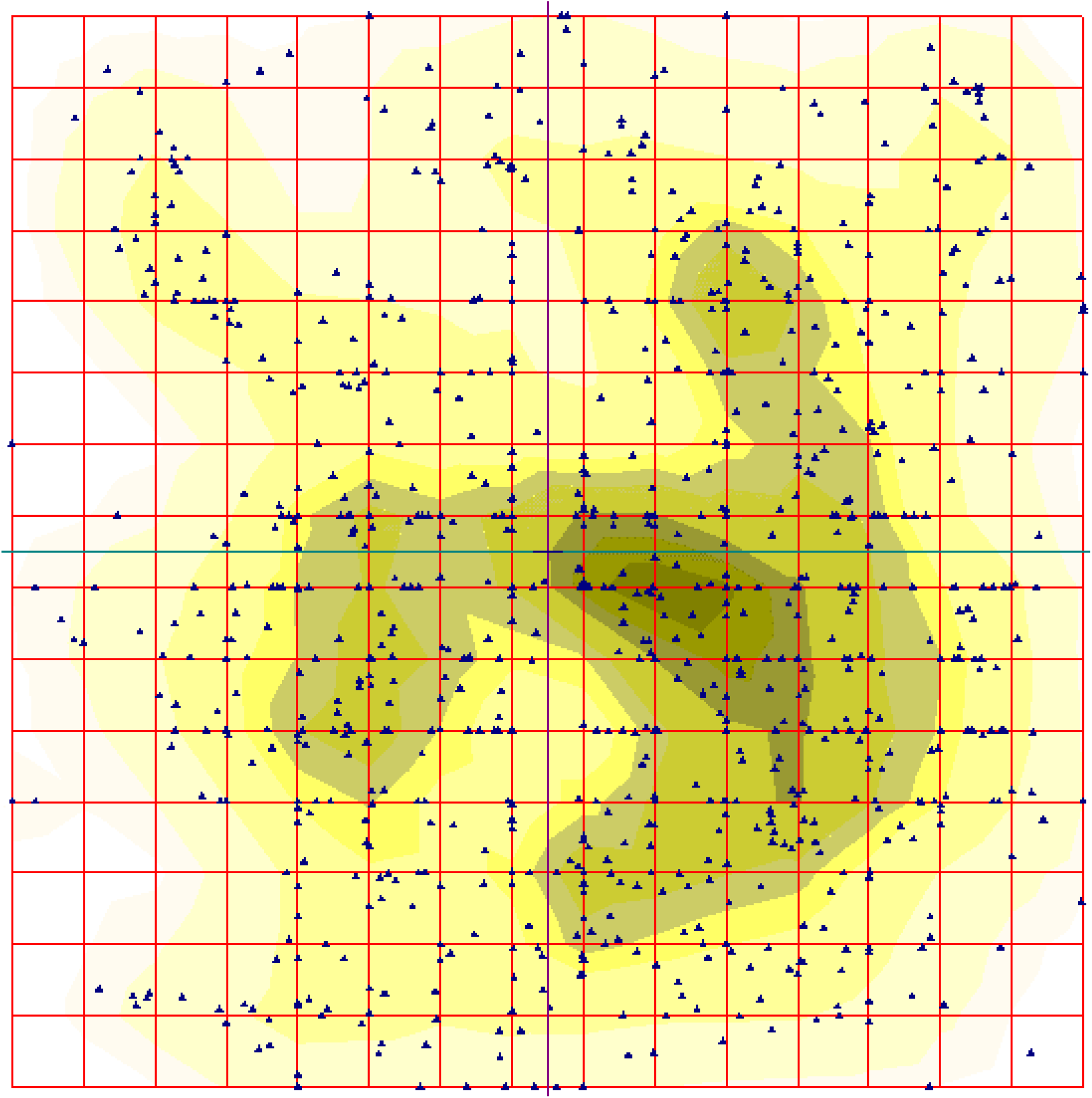
The distribution of the patients over the 16 × 16 elastic map.

**Fig 2.**
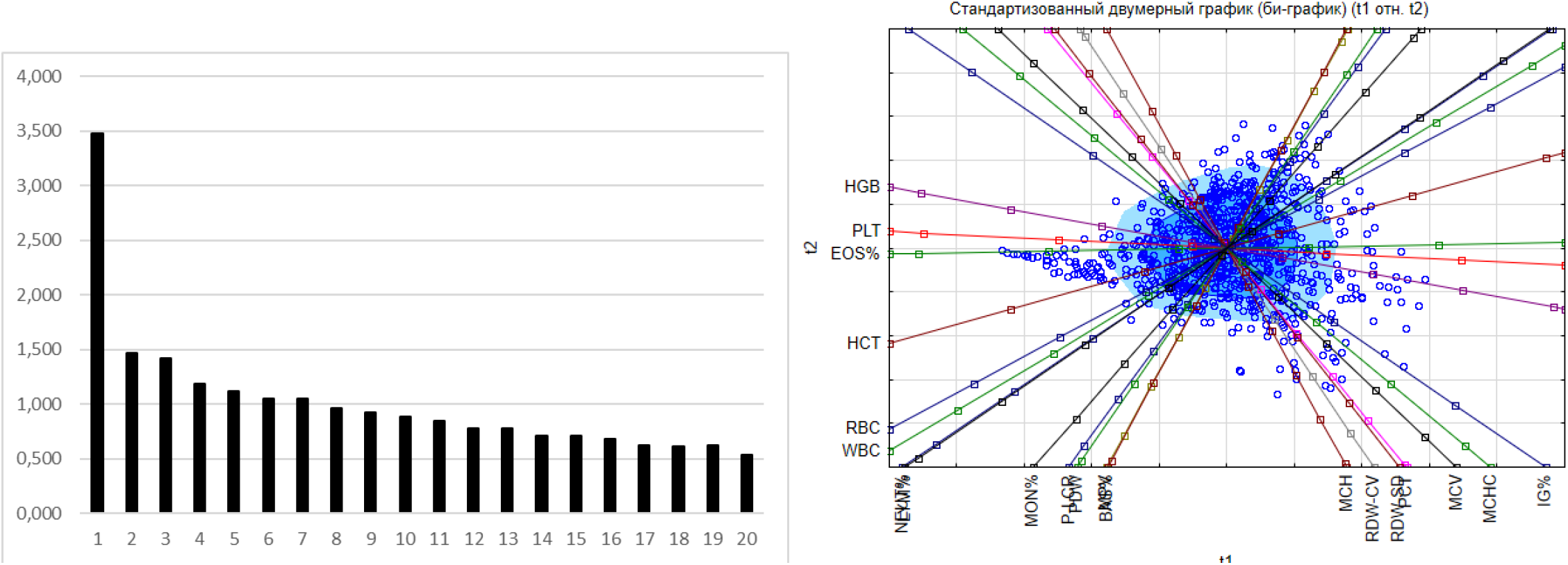
Eigenvalues variation for the data set under consideration.

Also, we studied the distribution of various types of tumors over the map; the types are shown in Table 1. To do it, the points on the map were labeled according to the localization or the type of tumor. A specific prevalence in a cluster occupation has been observed neither for localization, nor for the type (malignant or benign one) of tumor. Such effect may result from the significant bias in the tumor type distribution, in the original database: it comprises 720 records with malignant tumor diagnosis, and only 44 entries with benign ones. Similar absence of a prevalence in the distribution has been observed for sex or age of the patients.

At the next stage, we studied the distribution of the patients over the elastic map in dependence on the specific value of the used CBC parameters. Actually, the lowest and the highest values for each 21 parameter of CBC over the database could be found. It should be stressed, that some patients exhibit the figures falling beyond the normal span of that former, some do not. So, the span was split into 10 equal intervals, and we traced on the map the patients with different values of CBC characteristics. Step by step, we labeled the patients with the level of a character not exceeding λ*_j_*; here λ denotes a character, and j denotes the interval number, 1 ≤ *j* ≤ 10.

Surprisingly, it was found that all 21 character is divided into two groups, in terms of the pattern distribution of the patients over the elastic map, for increasing values of those characters. The pattern called *starry sky* was observed for BAS, EOS, WBC and IG. This pattern manifests in considerably homogeneous occurrence of the points on the elastic map, as the level of a character of CBC grows up. Surely, there are some minor variations in the details of the *starry sky* appearance, for different characters of CBC. Fig. 4 shows this starry sky type pattern observed for eosinophils content distribution over the map; these are eosinophils.

The opposite type of the distribution of the points over the elastic map called wave was observed for greater part of the characters of CBC. As soon, as a character (from this list) grows up from the least figure to the maximal one, points corresponding to the patients tend to occupy the map quite regularly, layer by layer. The pattern looks like a moving wave, indeed. Evidently, various characters have different starting locations on the map: it means that there are (almost) nobody at the database who has the minimal figures of two characters, simultaneously (see detailed discussion below). Fig. 3 shows this wave-type pattern observed for monocyte content distribution over the map.

**Fig 3.**
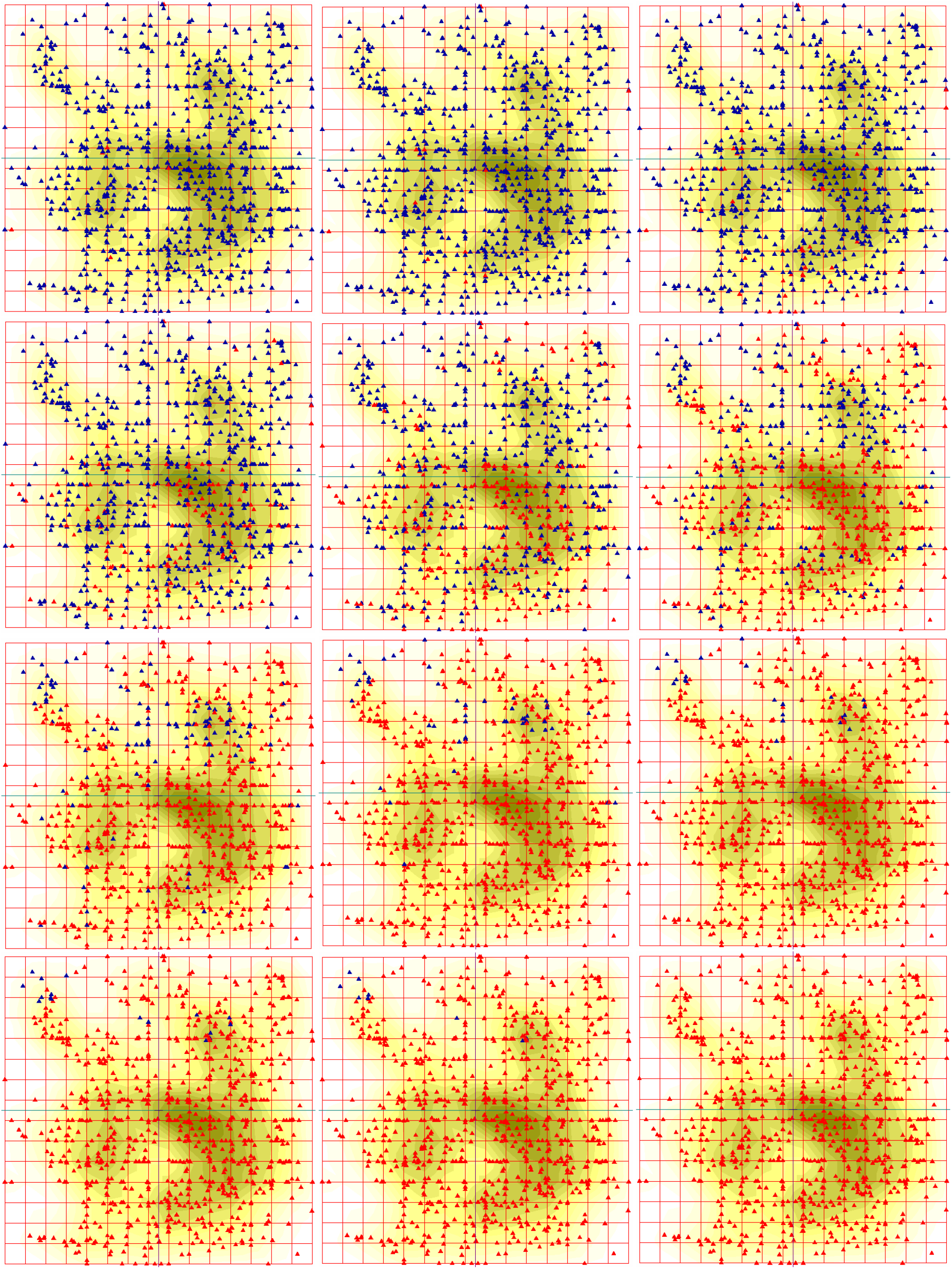
Wave-type distribution of the patients with various level of monocyte content over the 16 × 16 elastic map; see explanation in the text; monocytes.

**Fig 4.**
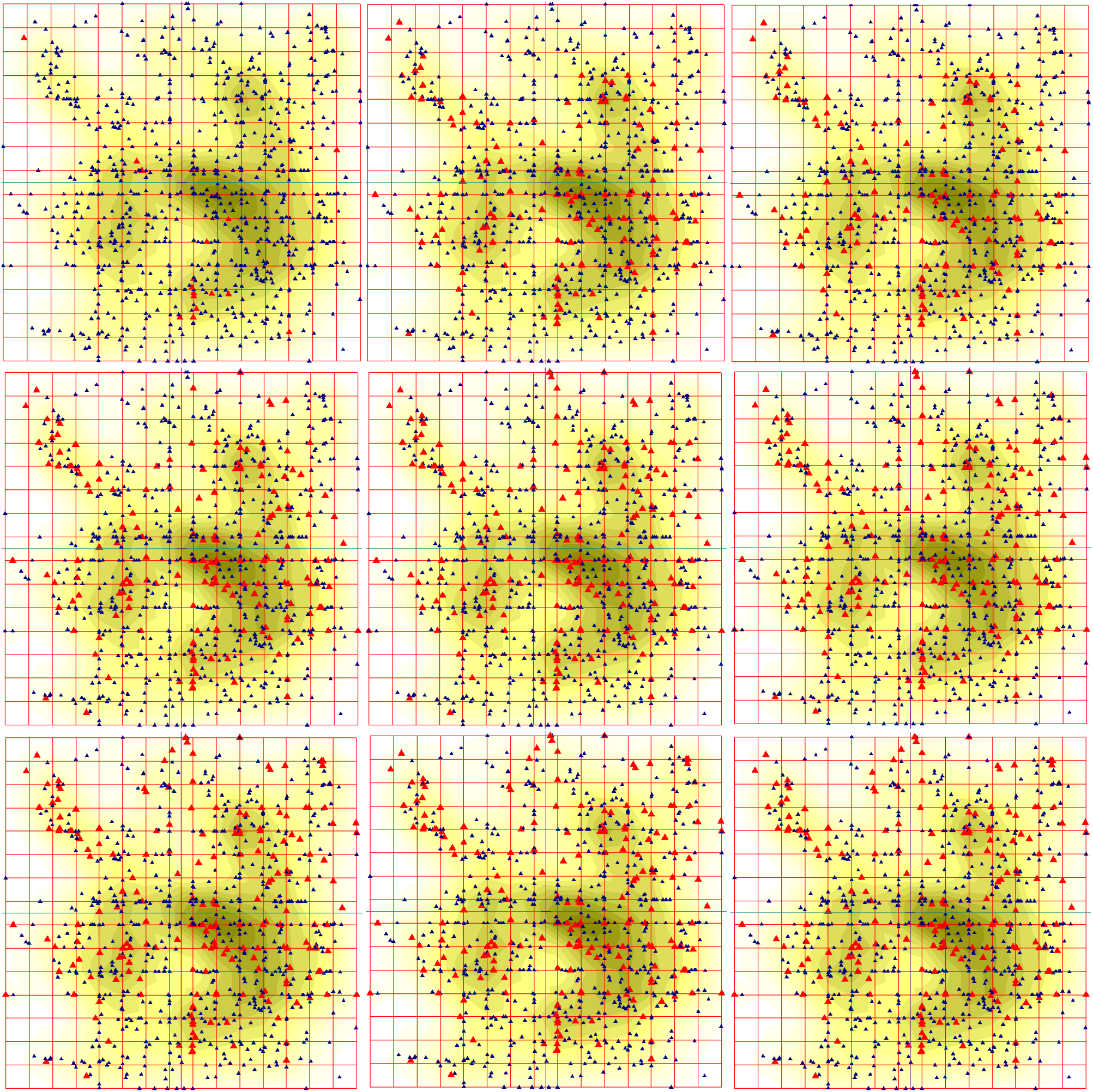
Starry-sky type distribution of the patients with various level of monocyte content over the 16 × 16 elastic map; see explanation in the text.

## Discussion

The occurrence of malignant neoplasms is a complicated process involving a number of factors. Currently, there is no universal and indisputable theory to classify these factors [22–25]. A number of paper brings an evidence of the extreme variety and diversity both of cancer tumors, and factors standing behind. Thus, the elabouration of logically apparent and informative predictors for malignant neoplasm development is still an acute problem, and sounding progress is reported in this direction [1, 4, 8, 18, 25]. Sophisticated molecular biology techniques stand behind this progress, mainly.

An implementation of the up-to-date advanced methods of nonlinear statistical analysis brings an opportunity to reveal knowledge from classical standard procedures and data records. Some methods and approaches may look quite complicated and unconventional thus requiring from a researcher some special efforts and qualification in data processing; very low cost, availability and routine standard procedures for the data records gathering introduced in every day practice of any hospital is the pay-off for these calamities.

There are two fundamental issues in the modern approach in medical data treatment based on big data methodology:

- the simultaneous and combined analysis of a number of different characteristics which originally had not been installed into a common theory is the first issue, and
- “modelling” of data, that is the approximation of multidimensional bulky data with a manifold of low dimension is the second issue.

Here we pursued both these ideas. Let’s focus on Table 1 once again. Neither elastic map technique (see Fig. 1), nor *K*-means identified a relation between a cluster, and a type of malignant neoplasm; same is true for benign neoplasms. Such failure may result from excessive specification in the localization and cancer type. It might be, gathering the different tumors having the same ontogenetic origin, one gets more clear and apparent correlation between clustering and the disease.

The features of the data base mentioned above make some constraints on the analysis and experiment design (if any). The impossibility to gather a reference group is the key point here, unless the specific factor is identified to arrange a randomization against that former. In practice, it means that there is no way to collect a group of healthy blood donors to be a reference group for further comparison with the patients with malignant neoplasms. The main reason of that failure is that comes from a number factors falling beyond any control, in a tentative reference group. For example, one is never able to provide a similar geography distribution of the reference group members, as well as occupation, etc. Meanwhile, all these factors may affect significantly the distribution pattern. Moreover, one has to gather a separate reference group for each specific malignant neoplasm included into the database. To overcome the problem, we analyzed the patients cohorts. The identified features of tumors with different localization may provide a good tool for alternative diagnostics and patient investigation.

Let now focus on Fig. 3; we have examined all the characteristics of CBC, in terms of a pattern of the map filling. It was found, various characteristics start to fill the map from different places, and do it in a specific way. This fact seems to be a manifestation of Liebig principle: any cancer tumor is a severe stress factor. Thus, the stress causes a mobilization of the resources of a sick organism concentrating them into a kind of a “bottle neck”. The most amazing fact here is that the data base contains a number of patients with **different** pathologies in **different** organs and tissues. And regardless the specific type of localization of a tumor, we see through the clustering provided by elastic map technique that a kind of specialization in the pathways and contours for the resources mobilization takes place.

Indeed, there is only one pair of variables that yields a pretty close pattern of filling out of the map, as the values grow up: these are haematocrit and haemoglobin. This situation seems to be quite natural: the correlation coefficient between these two variables is very high. These two CBC characteristics start to fill the map from the left border and go smoothly to the right keeping quite a straight line. On the contrary, erythrocytes start to fill the map from the upper left corner and do it as a arc wave centered at the left upper square vortex. Surprisingly, SER figures resembles to some extent the pattern of filling out observed for erythrocytes, while it takes start from the right lower corner of the square, thus completely opposing to erythrocytes.

Monocyte filling pattern looks like a straight front starting at the bottom of the map and going up smoothly, with minor variation of the front. Leucocytes start to fill the map from the upper right corner so that the starting distribution looks like a strap. That latter expands down and (slightly) left so that the left side of the map is filled when the highest values of monocyte content is observed.

Lymphocytes and average erythrocyte volume yield the most amazing filling patterns. They start to fill the map from two and three sites, respectively. Lymphocytes start to fill the map from the lower left and upper right corners. Surprisingly, the patients with the highest figures of the lymphocyte content are aggregated in the central part of the map occupying the area located along the “main” diagonal of that latter spreading from the left upper corner to the right lower one. The pattern of average erythrocyte volume distribution is unique it starts to fill the map from three sites. These latter are located in the left upper corner, right upper and left down ones.

The patterns of the map filling described above unambiguously prove the stratification of the patients over the space determined by CBC characteristics. There is no stratification of the patients in terms of the correspondence of the clusters identified by elastic map technique and age, sex or disease description (type and/or localization of a tumor). On the contrary, the patients are stratified in their respond type to cancer tumor attack. There is rather evident correlation between cluster composition, and the pattern of the map filling. That latter reflects the type and way of mobilization of the resources of an organism caused by malignant neoplasm, and the stratification strategy follows Liebig principle. This is basically new stratification observed in cancer patients, and further studies on the detailed relations between the respond type and related essential clinical aspects still are required.

## Acknowledgments

This study was supported be the grant from Voino-Yasenetsky Krasnoyarsk State Medical University, grant # 2.3 under the contract # 203.

## References

1. Arem H, Loftfield E. Cancer epidemiology: A survey of modifiable risk factors for prevention and survivorship. American journal of lifestyle medicine. 2018;12(3):200–210.

2. Kasting ML, Giuliano AR, Reich RR, Roetzheim RG, Nelson DR, Shenkman E, et al. Hepatitis C virus screening trends: serial cross-sectional analysis of the National Health Interview Survey Population, 2013-2015. Cancer Epidemiology and Prevention Biomarkers. 2018;27(4):503–513.

3. Albeshan SM, Mackey MG, Hossain SZ, Alfuraih AA, Brennan PC. Breast cancer epidemiology in gulf cooperation council countries: a regional and international comparison. Clinical breast cancer. 2018;18(3):e381–e392.

4. Almugren N, Alshamlan H. A Survey on Hybrid Feature Selection Methods in Microarray Gene Expression Data for Cancer Classification. IEEE Access. 2019;7:78533–78548.

5. Koval A, Katanaev VL. Dramatic dysbalancing of the Wnt pathway in breast cancers. Scientific reports. 2018;8(1):7329.

6. Blagodatski A, Cherepanov V, Koval A, Kharlamenko VI, Khotimchenko YS, Katanaev VL. High-throughput targeted screening in triple-negative breast cancer cells identifies Wnt-inhibiting activities in Pacific brittle stars. Scientific reports. 2017;7(1):11964.

7. Ahmed K, Koval A, Xu J, Bodmer A, Katanaev VL. Towards the first targeted therapy for triple-negative breast cancer: Repositioning of clofazimine as a chemotherapy-compatible selective Wnt pathway inhibitor. Cancer letters. 2019;449:45–55.

8. Dubey AK, Gupta U, Jain S. Breast cancer statistics and prediction methodology: a systematic review and analysis. Asian Pac J Cancer Prev. 2015;16(10):4237–4245.

9. Orlovskaya L, Vs O. nazvanie. Izvestiya VUZov, sev-kav TV. 2005;5(10):68–74.

10. Dolgova E. Sposob etc. Clinical Labouratory Diagnositcs. 2015;60(10).

11. Satparova L, Kogina E, Satparov Y, Galimov S, Knyazeva O, Udut V. Evaluation of some charateters of biochemical, clinical and immunological analysis of blood of the patients with mammalian cancer. In: Contemporary aspects of control and regulation; 2018. p. 246–253.

12. Arlekya I, Pleten’ A. Differential diagnostics of carcinoid tumors of gastrointestinal tract. In: Fundametal problems of science; 2017. p. 131–134.

13. Taş EE, Özgen E, Yavuz AF. Can Preoperative Complete Blood Count Parameters Be Used as Predictive Markers for Lymph Node Metastasis in Endometrial Carcinomas? Cyprus Journal of Medical Sciences. 2018;3(3):168–172.

14. Yayla Abide C, Bostanci Ergen E, Cogendez E, Kilicci C, Uzun F, Ozkaya E, et al. Evaluation of complete blood count parameters to predict endometrial cancer. Journal of clinical laboratory analysis. 2018;32(6):e22438.

15. Hornbrook MC, Goshen R, Choman E, O’Keeffe-Rosetti M, Kinar Y, Liles EG, et al. Correction to: Early Colorectal Cancer Detected by Machine Learning Model Using Gender, Age, and Complete Blood Count Data. Digestive diseases and sciences. 2018;63(1):270–270.

16. Esteva A, Kuprel B, Novoa RA, Ko J, Swetter SM, Blau HM, et al. Dermatologist-level classification of skin cancer with deep neural networks. Nature. 2017;542(7639):115.

17. Ogasawara A, Matsushita H, Tanaka Y, Shirasugi Y, Ando K, Asai S, et al. A simple screening method for the diagnosis of chronic myeloid leukemia using the parameters of a complete blood count and differentials. Clinica Chimica Acta. 2019;489:249–253.

18. Kourou K, Exarchos TP, Exarchos KP, Karamouzis MV, Fotiadis DI. Machine learning applications in cancer prognosis and prediction. Computational and structural biotechnology journal. 2015;13:8–17.

19. Fukunaga K. Introduction to statistical pattern recognition. London: Academic Press; 1990.

20. Gorban AN, Zinovyev AY. Fast and user-friendly non-linear principal manifold learning by method of elastic maps. In: 2015 IEEE International Conference on Data Science and Advanced Analytics, DSAA 2015, Campus des Cordeliers, Paris, France, October 19-21, 2015; 2015. p. 1–9. Available from: https://doi.org/10.1109/DSAA.2015.7344818.

21. Xu D, Tian Y. A Comprehensive Survey of Clustering Algorithms. Annals of Data Science. 2015;2(2):165–193. doi:10.1007/s40745-015-0040-1.

22. Brown KF, Rumgay H, Dunlop C, Ryan M, Quartly F, Cox A, et al. The fraction of cancer attributable to modifiable risk factors in England, Wales, Scotland, Northern Ireland, and the United Kingdom in 2015. British journal of cancer. 2018;118(8):1130.

23. Sakamoto Y, Kokudo N, Matsuyama Y, Sakamoto M, Izumi N, Kadoya M, et al. Proposal of a new staging system for intrahepatic cholangiocarcinoma: analysis of surgical patients from a nationwide survey of the Liver Cancer Study Group of Japan. Cancer. 2016;122(1):61–70.

24. Islami F, Goding Sauer A, Miller KD, Siegel RL, Fedewa SA, Jacobs EJ, et al. Proportion and number of cancer cases and deaths attributable to potentially modifiable risk factors in the United States. CA: a cancer journal for clinicians. 2018;68(1):31–54.

25. Dubey AK, Gupta U, Jain S. Epidemiology of lung cancer and approaches for its prediction: a systematic review and analysis. Chinese journal of cancer. 2016;35(1):71.

